# Proton-pumping rhodopsins in marine diatoms

**DOI:** 10.1101/2022.01.18.476826

**Authors:** Susumu Yoshizawa, Tomonori Azuma, Keiichi Kojima, Keisuke Inomura, Masumi Hasegawa, Yosuke Nishimura, Masuzu Kikuchi, Gabrielle Armin, Hideaki Miyashita, Kentaro Ifuku, Takashi Yamano, Adrian Marchetti, Hideya Fukuzawa, Yuki Sudo, Ryoma Kamikawa

**Affiliations:** Atmosphere and Ocean Research Institute, The University of Tokyo, Chiba 277–8564, Japan; Graduate School of Frontier Sciences, The University of Tokyo, Chiba 277–8563, Japan; Collaborative Research Institute for Innovative Microbiology, The University of Tokyo, Tokyo 113–8657, Japan; Graduate School of Human and Environmental Studies, Kyoto University, Kyoto 606-8501, Japan; Division of Pharmaceutical Sciences, Okayama University, Okayama 700-8530, Japan; Graduate School of Medicine, Dentistry and Pharmaceutical Sciences, Okayama University, Okayama 700-8530, Japan; Graduate School of Oceanography, University of Rhode Island, Narragansett, RI, USA; Graduate School of Agriculture, Kyoto University, Kyoto 606-8502, Japan; Graduate School of Biostudies, Kyoto University, Kyoto, 606-8502, Japan; Department of Earth, Marine and Environmental Sciences, University of North Carolina at Chapel Hill, Chapel Hill, North Carolina, USA

## Abstract

Diatoms are a major phytoplankton group responsible for about 20% of Earth’s primary production. They carry out photosynthesis inside the plastid, an organelle obtained through eukaryote-eukaryote endosymbiosis. Recently, microbial rhodopsin, a photoreceptor distinct from chlorophyll-based photosystems, has been identified in certain diatoms. However, the physiological function of diatom rhodopsin is not well understood. Here we show that the diatom rhodopsin acts as a light-driven proton pump and localizes to the outermost membrane of the four membrane-bound complex plastids. Heterologous expression techniques were used to investigate the protein function and subcellular localization of diatom rhodopsin. Using model simulations, we further evaluated the physiological role of the acidic pool in the plastid produced by proton-transporting rhodopsin. Our results propose that the rhodopsin-derived acidic pool may be involved in a photosynthetic CO_2_-concentrating mechanism and assist CO_2_ fixation in diatom cells.

## Introduction

Diatoms are unicellular, photosynthetic algae found throughout aquatic environments, and are responsible for up to 20% of annual net global carbon fixation^1^. As their contribution to primary production in the ocean is significant, their light utilization mechanisms are essential to correctly understand marine ecosystems. Diatoms contain chlorophyll a and c and carotenoids such as fucoxanthin as photosynthetic pigments in the secondary plastids acquired by eukaryote-eukaryote endosymbiosis^2^. In addition, certain diatoms have recently been shown to contain microbial rhodopsin (henceforth rhodopsins), a light-harvesting antenna distinct from the chlorophyll-containing antenna for photosynthesis^3^. Although rhodopsin-mediated light-harvesting may somehow support the survival of these diatoms in marine environments, the physiological role of rhodopsin in diatom cells remains unclear.

Microbial rhodopsins are a large family of seven transmembrane photoreceptor proteins^4^. Rhodopsin has an all-*trans* retinal as the light-absorbing chromophore, and its protein function is triggered by light-induced isomerization of the retinal. The first microbial rhodopsin, the light-driven proton pump bacteriorhodopsin (BR), was discovered in halophilic archaea^5^. Although rhodopsin was initially thought to occur only in halophilic archaea inhabiting hypersaline environments, subsequent studies have shown that the rhodopsin gene is widely distributed in all three domains of life^6,7^. Rhodopsins can be classified based on their functions into light-driven ion pumps, light-activated signal transducers, and light-gated ion channels. The two former functional types of rhodopsin have so far been found in prokaryotes^8^. The rhodopsins in prokaryotes, regardless of function, are localized to the plasma membrane in which they operate. For example, proton-pumping rhodopsins export protons from the cytosol across the plasma membrane to convert light energy into a proton motive force (PMF)^9^. The PMF induced by rhodopsin ion transport is utilized by various physiological functions such as ATP synthesis, substrate uptake, and flagellar movement.

In eukaryotic microorganisms, rhodopsins functioning as light-driven ion pumps and light-gated ion channels have been reported^8,10^. Light-gated ion channels are well studied; they exclusively localize in the eyespot of phytoplankton and are involved in phototaxis^11^. The other type of rhodopsin in eukaryotes, light-driven ion-pumping rhodopsins have been found in a number of organisms belonging to both photoautotrophic and heterotrophic protists^3,12^. Since the intracellular membrane structure of eukaryotic cells is more complex than that of prokaryotes, containing a variety of organelles, even light-driven ion-pumping rhodopsins may have distinct physiological roles depending on their subcellular localization^12^. However, due to the difficulty in determining the exact localization of rhodopsins in eukaryotic cells, most rhodopsins are not even known for their subcellular localization.

In this study, to clarify the physiological function of rhodopsin in a marine pennate diatom, we investigated the phylogeny, protein function, spectroscopic characteristics, and subcellular localization of rhodopsin from a member of the genus *Pseudo-nitzschia*. Heterologous expression techniques were used to analyze the protein function and spectroscopic features. The expression eGFP-fusion rhodopsin also revealed the subcellular localization in the model diatom (*Phaeodactylum tricornutum*). Furthermore, a model-based analysis was performed to evaluate the impacts of the potential roles of the rhodopsin for cellular biology.

## Results and Discussion

### Rhodopsin sequences and phylogenetic analysis

We performed phylogenetic analysis using the rhodopsin (named PngR, accession no. AJA37445.1) of the diatom *Pseudo-nitzschia granii* and the microbial rhodopsin sequences reported to date^3^. This phylogenetic tree revealed that PngR is not included in the proteorhodopsin (PR) clade commonly found in the ocean but belongs to the Xanthorhodopsin (XR) -like rhodopsin (XLR) clade, which is presumed to have an outward proton transporting function (Fig. 1 and Extended Data Fig. 1). The comparison of motif sequences necessary for ion transport showed that the amino acids in the putative proton donor and acceptor sites of XR and PR were conserved in PngR, suggesting that PngR functions as an outward proton pump (Extended Data Fig. 2). Furthermore, the homology search for rhodopsin sequence in the XLR clade from Marine Microbial Eukaryote Transcriptome Sequencing Project (MMETSP) revealed that not only diatoms (Ochrophyta, Stramenopiles), but also dinoflagellates (Dinophyceae, Alveolata) and haptophytes have rhodopsin genes in the same XLR clade (Supporting Information Table S1). These results indicate that rhodopsin of the XLR clade is widely distributed among the major phytoplankton groups, which are important primary producers in the ocean.

**Figure 1.**
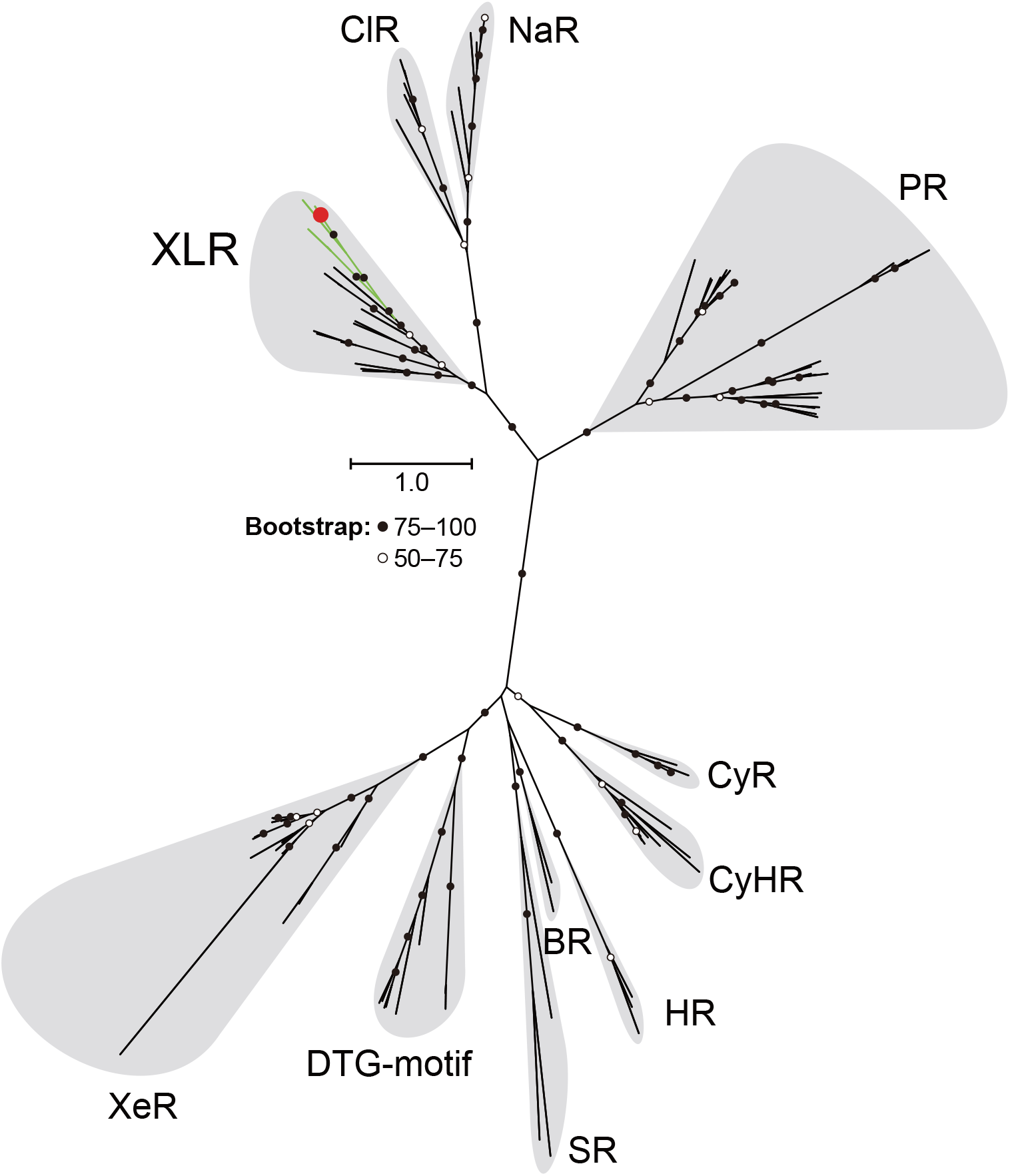
Phylogenetic position of diatom rhodopsin. Maximum likelihood tree of amino acid sequences of microbial rhodopsins. Diatom rhodopsin (PngR) is indicated by a red circle and bootstrap probabilities (≥ 50%) are indicated by black and white circles. Green branches indicate eukaryotic rhodopsins used in this analysis, and black branches indicate others. Rhodopsin clades are as follows: XLR (Xanthorhodopsin-like rhodopsin), ClR (Cl^-^-pumping rhodopsin), NaR (Na^+^-pumping rhodopsin), PR (proteorhodopsin), XeR (xenorhodopsin), DTG-motif rhodopsin, SR (sensory rhodopsin-I and sensory rhodopsin-II), BR (bacteriorhodopsin), HR (halorhodopsin), CyHR (cyanobacterial halorhodopsin), and CyR (cyanorhodopsin).

### Function and spectroscopic features of Diatom rhodopsin

To characterize the function of the PngR, we heterologously expressed the synthesized rhodopsin gene in *Escherichia coli* cells. A light-induced decrease in pH was observed in the suspension of the *E. coli* cells expressing the PngR, and this decrease was almost completely abolished in the presence of the protonophore carbonyl cyanide M-chlorophenylhydrazone (CCCP) (Fig. 2A). The observed pH changes clearly showed that PngR exports protons from the cytoplasmic side across the plasma membrane.

**Figure 2.**
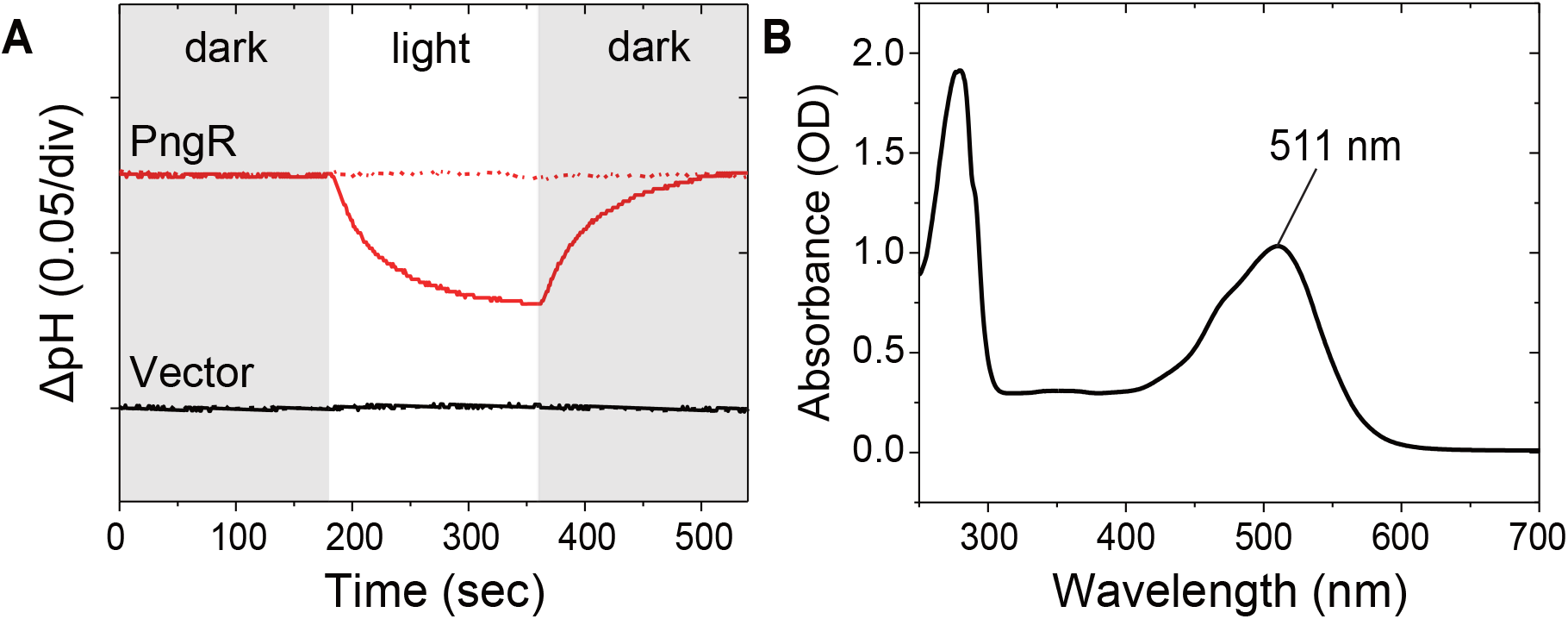
Light-induced pH changes and absorption spectrum of PngR. (A) Outward proton pump activity of PngR in *E. coli* cells. Light-induced pH changes of solutions containing *E. coli* cells with the expression plasmid for PngR (upper panel) and the empty vector pET21a (lower panel) in the presence (red dashed line) or absence (red solid line) of CCCP. The white-filled region indicates the period of illumination. (B) Absorption spectrum of purified PngR in Buffer A (50 mM Tris–HCl, pH 7.0, 1 M NaCl and 0.05% (w/v) DDM).

Next, we examined the spectroscopic characteristics of PngR using recombinant protein purified from *E. coli*. The absorption maximum of PngR was located at 511 nm (Fig. 2B), which was explicitly shorter than that of XR (565 nm) and GR (*Gloeobacter* rhodopsin 541nm) in the XLR clade^13^. It is noteworthy that while *P. granii* is a marine species, while XR and GR are both distributed in terrestrial freshwater organisms. Thus, our observations are consistent with the shorter wavelength of the absorption maximum of rhodopsin in marine environments than in the terrestrial environment^14^, indicating that PngR is well adapted to the light conditions in the ocean.

We then examined the retinal configuration in PngR by high-performance liquid chromatography. In both in light- and dark-adapted samples, the isomeric state of retinal was predominantly all-*trans* (Extended Data Fig. 3), which was similar to the isomeric state of retinal in prokaryotic GR in XLR clade but different from that in BR^15^. Since the p*K*_a_ value of the proton acceptor residue (Asp85 in BR) is an indicator of the efficiency of proton transport by rhodopsin, we estimated the p*K*_a_ values of the putative proton acceptor in PngR (Asp91) by pH titration experiment (Extended Data Fig. 4). This experiment estimated that the p*K*a of this residue acceptor is about 5.0, indicating that the proton acceptor of PngR works well in marine and intracellular environments. Furthermore, the photochemical reactions that proceed behind the ion-transportation mechanism of PngR were examined by flash-photolysis analysis (Extended Data Fig. 5). All photocycle of PngR were completed in about 300 ms, suggesting that the cycle is fast enough to transport protons in a physiologically significant time scale. Detailed results and discussion of flash-photolysis analysis are described in the supplementary information.

### Subcellular localization of PngR in a model diatom

The PngR sequence bears neither apparent N-terminal extension nor detectable N-terminal signal peptide, and thus *in silico* analyses are not able to predict a possible subcellular localization of PngR. To identify the subcellular localization of PngR, a C-terminal eGFP-fusion PngR was expressed in the model diatom *Phaeodactylum tricornutum*, which can be transformed by electroporation and is often used for heterologous expression analysis^16,17^. The transformed *P. tricornutum* cell was observed with differential interface contrast and epifluorescent microscopes (Fig. 3A). From these microscopic observations, we examined the morphology of *P. tricornutum* and the localization of recombinant PngR, nucleus, chlorophyll and other cellular organelles in multiple cells (Extended Data Fig. 6 and 7). The fluorescence signal of PngR:eGFP transformant was most likely localized in the periphery of chlorophyll fluorescence and DAPI signals, corresponding to the outermost plastid membrane, called the chloroplast endoplasmic reticulum membrane (CERM), which is physically connected to the nuclear membrane (Fig. 3A). A few cells also exhibited GFP signals within vacuolar membranes in addition to CERM (Extended Data Fig. 7). Insertion of the complete sequence of the PngR:eGFP gene in the transformant DNA was confirmed by PCR followed by the Sanger sequencing.

**Figure 3.**
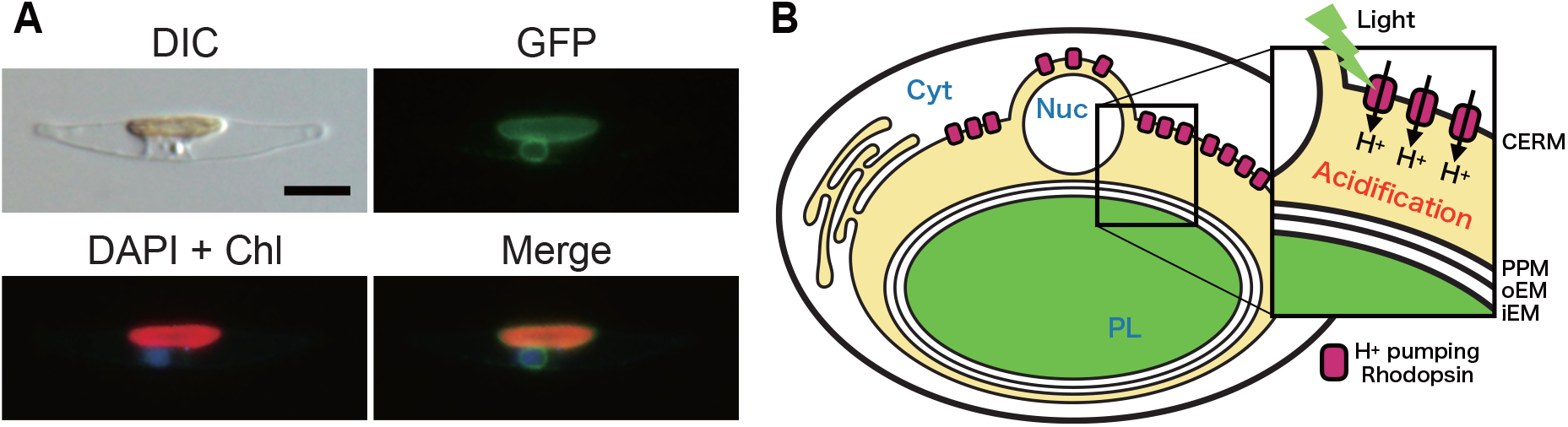
Subcellular localization of rhodopsins in diatom cell. (A) A transformed diatom cell was observed with DIC (Differential Interface Contrast) (Upper left). Green fluorescence from the recombinant PngR (GFP) (Upper right). The nuclear DNA stained with DAPI and the chlorophyll autofluorescence (DAPI + Chl) and merged image (Merge) are shown in bottom left and bottom right, respectively. Scale bar indicates 5 μm. (B) A mode for subcellular localization of PngR. The proton transport of PngR acidifies the region (the middle space) surrounded by the membrane of CERM and PPM. Abbreviations are as follows: Cyt (Cytosol), Nuc (Nucleus), PL (Plastid), CERM (Chloroplast endoplasmic reticulum membrane), PPM (Periplastidial membrane), oEM (Outer plastid envelop membrane) and iEM (Internal plastid envelop membrane).

Given the above heterologous expression experiment and microscopic observation, we expect that PngRs primarily localize to the outermost membrane of the plastid, although to a lesser extent, may also be found in the vacuolar membrane, which may be a function of other factors such as cell growth conditions. In addition, PngR should be oriented in the direction that the inside of the CERM is regarded as the outside of an analogous prokaryotic cell. These results imply that the light-driven proton transport by PngR could acidify the inner region of the CERM (Fig. 3B). If correct, what physiological role does the acidification of this region have in diatoms? The electrochemical gradient formed by rhodopsin could be a driving force for various secondary transport processes. Alternatively, since the primary purpose of plastids is photosynthesis, the formation of acidic pools is likely to be somehow related to photosynthesis. One possibility is that the appearance of this acidic pool influences carbon fixation, since in the carbonate system, acidification favors a higher concentration of CO_2_ relative to the other dominant carbonate species in seawater (HCO_3_^-^and CO_3_^2-^).

In general, under weakly alkaline conditions in the ocean, most of the dissolved inorganic carbon (DIC) is present in the form of HCO_3_^-^, and only approximately 1% is present in the form of CO_2_. However, ribulose-1,5-bisphosphate carboxylase/oxygenase (RuBisCO) localized in the stroma can only react with inorganic carbon (Ci) in the form of CO_2_, but not of HCO_3_^-^. The RuBisCO enzyme in diatoms has a low affinity even for CO_2_ (Km of 25∼68 μM while CO_2_aq in the ocean is about 10 μM at 25°C) and thus requires concentrated CO_2_ for efficient fixation at the site of RuBisCO. In other words, the ocean has always been a CO_2_-limited environment for most phytoplankton^18^. Consequently, due to membrane impermeability of HCO_3_^-^, phytoplankton have developed a variety of CO_2_-concentrating mechanism (CCM) to efficiently transport Ci to the site of RuBisCO by placing HCO_3_^-^transporters in appropriate membranes and carbonic anhydrase (CA) in these compartments, the latter of which catalyzes the rapid interconversion between HCO_3_^-^and CO_2_. This interconversion can be facilitated if the balance of the carbonate system is biased toward CO_2_ with acidification.

In marine phytoplankton, especially those with secondary plastids, since RuBisCO is located in the stroma surrounded by three or four membranes, it is necessary to localize HCO_3_^-^transporters in each membrane^19^. Diatoms that possess four membrane-bound plastids are known to have a HCO_3_^-^transporter (e.g., SLC4) present in the plasma membrane^16^, but it is not well understood how CO_2_ is concentrated in close proximity to RuBisCO in the stroma^20^. In sharp contrast to the transporter-based mechanism, if the equilibrium of Ci can be shifted toward CO_2_ by acidifying the inner region of the outermost membrane, it may be possible to transport CO_2_ to RuBisCO in a membrane-permeable form through gas diffusion. However, since it is difficult to directly investigate the role of membrane protein in eukaryotic microbial organelles, the combination of observation and model simulation is a powerful approach^21^.

### A quantitative model of Carbon concentration in diatoms: CFM-CC

Our subcellular localization analysis suggests that proton transport by rhodopsin acidifies the inner side of the outermost membrane of the plastid (hereafter “the middle space”). To quantitatively test the effect of the decreased pH in the middle space on C fixation, we developed a simple quantitative model of carbonate chemistry combined with membrane transport and C fixation (CFM-CC: Cell Flux Model of C Concentration) (Fig. 4 upper panel). Previously, a comprehensive model of CO_2_ concentration within diatoms was developed^22^. CFM-CC uses a conceptually similar structure to this model, focusing on more specific membrane layers, designed to test the effect of altered pH in the middle space.

**Figure 4.**
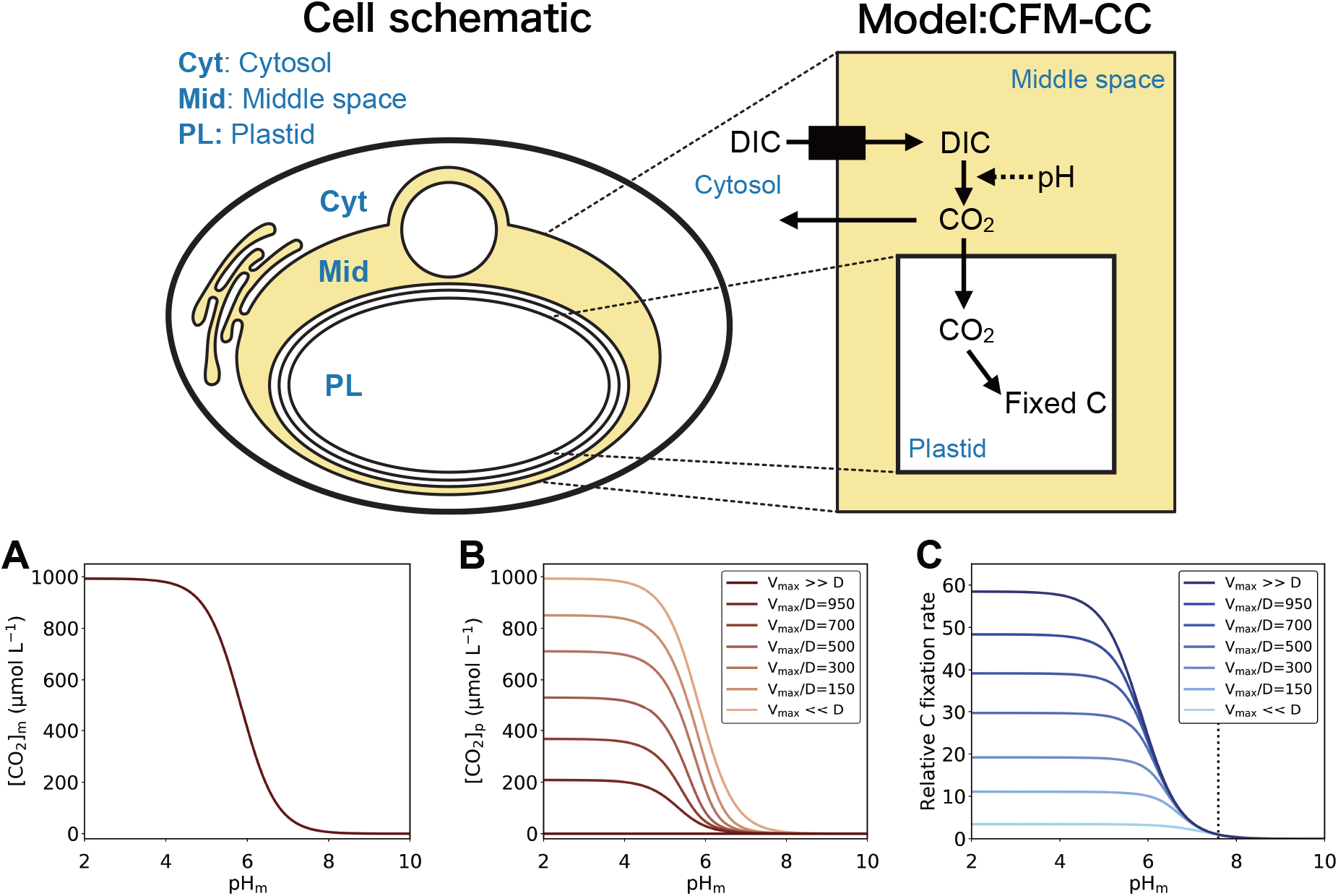
A quantitative model of carbon concentration in diatoms. (Upper panel) Schematic of Cell Flux Model of C Concentration (CFM-CC). The left panel represents the actual cell and the right panel represents the model. Solid arrows show net flux of C and dashed arrow indicates the influence of pH. (Bottom panel) The influence of pH in the middle space on CO_2_ concentrations and the photosynthesis rate. (A) CO_2_ concentrations in the middle space [*CO*_2_]_m_. (B) CO_2_ concentrations in the plastid [*CO*_2_]_p_.(C) C fixation rate relative to that with pH in the middle space of 7.59, the only value we found for intracellular pH in a diatom^23^. (B) and (C) are plotted for various *V*_*max*_*/D*. The solution for [CO_2_]_m_ in (A) is independent of *Vmax/D*.

Our model result shows that the concentrations of CO_2_ in the middle space ([*CO*_2_]_*m*_) strongly depend on the pH (pH_m_), suggesting that proton pumping by rhodopsin could enhance C fixation (Fig. 4 bottom panel). The calculation of C chemistry in the middle space shows that decreasing pH_m_ favors higher [*CO*_2_]_*m*_ at a given DIC concentration (Fig. 4A) (here we used 993 μmol L^-1 23^). We note that the potential leaking of CO_2_ to the cytosol may change the DIC in the middle space, but here we use a constant DIC value, since such effect has not been experimentally shown and is difficult to quantify due to unknown factors (e.g., the balance of DIC uptake and CO_2_ leaking). At the reference point (here pH_m_ = 7.59 ^23^), [*CO*_2_]_*m*_ is 17 μmol L^-1^ [y007 03], but it increases to 64, 410, and 870 μmol L^-1^ for pH_m_ values of 7, 6, and 5, respectively [y007 04] (Fig. 4A).

Due to the increased [*CO*_2_]_*m*_, CO_2_ concentration in the plastid [*CO*_2_]*p* also increases with lowered pH_m_, but the level of increase depends on *V*_*max*_*/D* (the ratio of maximum C fixation rate to the diffusion constant) (Fig. 4B). When *V*_*max*_*/D* is small, diffusion of CO_2_ from the middle space to the plastid dominates the influence, resulting in [*CO*_2_]_*p*_ similar to [*CO*_*2*_]_*m*_. In contrast, when *V*_*max*_*/D* is large, CO_2_ uptake dominates, and there is small influence of [*CO*_2_]_*m*_ on [*CO*_2_]_*p*_.

The rate of C fixation increases with lowered pH_m_, since the increased [*CO*_2_]_*m*_ leads to faster transport of CO_2_ into the plastid (Fig. 4C). However, the magnitude of such increase depends on *V*_*max*_*/D*. The model shows that the effect of pH_m_ on C fixation is larger when *V*_*max*_*/D* is large, because the rate of C fixation is more strongly affected by [*CO*_2_]_*m*_ [eq. 6], which is directly influenced by pH_m_. However, when *V*_*max*_*/D* is small, the C fixation rate is more strongly influenced by the C uptake kinetics [eq. 4], which saturates at relatively low [*CO*_2_]_*p*_ (*K* value of 44 μmol L^-1 24^). The increase in C fixation ranges 2.1-3.8, 3.2-24.2 and 3.4-51.2 times when pH_m_ decreases from 7.59 to 7, 6, 5 respectively [y008 13]. These results suggest two things. First, any *V*_*max*_*/D*, pH_m_ has a considerable effect on C uptake. Second, it would be highly favorable for the cells to have high *V*_max_, relative to the diffusivity of CO_2_ across the membrane. This is most likely the case since when there are multiple membranes between the middle space and plastid, which would lower the *D* value ^25,26^. Thus, this simple yet elegant system with rhodopsin to manipulate the pH_m_ can provide a powerful mechanism in C concentration and thus support a high C fixation in some rhodopsin-containing diatoms, enabling them to be more productive phytoplankton in the ocean.

Such a CCM based on CO_2_ diffusion (termed pump-leak type) is known to be a possible mechanism by placing CAs in appropriate locations. For example, *Nannochloropsis oceanica* (*Ochrophyta*), possessing the same four membrane-bound complex plastids as found in diatoms, is thought to generate CO_2_ by placing CA in the middle space^27^. Besides, centric diatom *Chaetoceros gracilis* is thought to use externally placed CAs to allow CO_2_ to flow into the cell^28^. In contrast to the CA-based model, acidification-based model was formerly proposed to facilitate CO_2_ fixation of RuBisCO in the thylakoid lumen of plastids; HCO_3_^-^is converted into CO_2_ by acidification by photosynthetic proton pumping into the thylakoid lumen^29^. Our model further expands the model and proposes that the middle space acidified by proton-pumping rhodopsin also plays the role of a CO_2_ generator. Our proposed mechanism would be useful in most parts of the ocean where CO_2_ chronically limits photosynthesis, but would be even more valuable in specific environments. For example, since CA, which plays a central role in CCM, requires cobalt or zinc ions as the reaction center, and photosynthetic proton pumping systems require iron, rhodopsin-derived acidic pools may be useful for the transport of Ci in oceans where these metal ions are depleted (such as the HNLC region of the North Pacific Ocean)^30^. Indeed, in the HNLC region of the North Pacific, where the *P. granii* rhodopsin-containing cells were initially isolated and sequenced^31^, it has been pointed out that primary production may be limited by iron and influenced by other trace metals^32^. In other words, this mechanism seems to be particularly effective in the ocean, where trace metals involved in CCM are depleted.

## Conclusion

In this study, we have clarified the function and subcellular localization of PngR in a photosynthetic diatom. Our results suggested that proton transport by rhodopsin creates a local acidic pool inside the outermost membrane of the plastid. Based on the quantitative simulation, we propose a conceptual model of rhodopsin-contributing CCM (Extended Data Fig. 8). The acidic pool created by light is likely involved in the CCM and provides positive feedback on carbon fixation efficiency. Future analyses of cultured rhodopsin-bearing microbial eukaryotes would corroborate our results and allow for the evaluation of how rhodopsin-mediated proton transport promote their growth in the ocean.

## Methods

### Rhodopsin sequences and phylogenetic analysis

The rhodopsin sequence of the diatom *Pseudo-nitzschia granii* was previously reported^3^. All other rhodopsin sequences used in the phylogenetic analysis were collected from the National Center for Biotechnology Information. Detailed information on the strains used in this analysis is given in Extended Data Fig. 1. The sequences were aligned using MAFFT version 7.453 with options’--genafpair’ and ‘--maxiterate 1000’. The phylogenetic tree was inferred using RAxML (v.8.2.12) with the ‘PROTGAMMALGF’ model using 1000 rapid bootstrap searches. Model selection was performed with the ProteinModelSelection.pl script in the RAxML package.

The search for eukaryotic rhodopsins that belong to the XLR clade was performed among protein sequences of Marine Microbial Eukaryote Transcriptome Sequencing Project (MMETSP)^33^. Namely, the phylogenetic placement of rhodopsin proteins from MMETSP using pplacer (v1.1.alpha19)^34^ was performed on a prebuilt large-scale phylogenetic tree of rhodopsins and extracted placements on the XLR clade using gappa (v0.6.0)^35^.

### Gene preparation, protein expression and ion transport measurements of *E. coli* cells

The full-length cDNA for PngR, whose codons were optimized for *E. coli*, were chemically synthesized by Eurofins Genomics and inserted into the NdeI-XhoI site of the pET21a(+) vector as previously described^36^. A hexa-histidine-tag was fused at the C-terminus of PngR, which was utilized for purification of the expressed protein. The heterologous protein expression method is the same as previously reported^37^. *E. coli* BL21(DE3) cells harboring the cognate plasmid were grown at 37 °C in LB medium supplemented with ampicillin (final concentration = 50 μg mL^-1^). Protein expression was induced at an optical density at 600 nm of 0.7–1.2 with 1 mM isopropyl β-d-1-thiogalactopyranoside (IPTG) and 10 μM all-trans retinal, after which the cells were incubated at 37 °C for 3 h. The proton transport activity of PngR was measured as light-induced pH changes of suspensions of *E. coli* cells as previously described^37^. In short, cells expressing PngR were washed more than three times in 150 mM NaCl and were then resuspended in the same solution for measurements. Each cell suspension was placed in the dark for several min and then illuminated using a 300 W Xenon lamp (ca. 30 mW cm^-2^, MAX-303, Asahi spectra, Japan) through a > 460 nm long-pass filter (Y48, HOYA, Japan) for 3 min. Measurements were repeated under the same conditions after the addition of the protonophore carbonyl cyanide m-chlorophenylhydrazone (CCCP) (final concentration = 10 μM). Light-induced pH changes were monitored using a Horiba F-72 pH meter. All measurements were conducted at 25 °C using a thermostat (Eyela NCB-1200, Tokyo Rikakikai Co. Ltd, Japan).

### Purification of PngR from *E. coli* cells and spectroscopic measurements of the purified rhodopsin

*E. coli* cells expressing PngR were disrupted by sonication for 30 min in ice-cold water in a buffer containing 50 mM Tris–HCl (pH 7.0) and 300 mM NaCl. The crude membrane fraction was collected by ultracentrifugation and solubilized with 1.0% (w/v) n-dodecyl-β-d-maltoside (DDM, DOJINDO Laboratories, Japan). The solubilized fraction was purified by Ni^2+^ affinity column chromatography with a linear gradient of imidazole as described previously^38^. The purified protein was concentrated by centrifugation using an Amicon Ultra filter (30,000 M_w_ cut-off; Millipore, USA). The sample media was then replaced with Buffer A (50 mM Tris–HCl, pH 7.0, 1 M NaCl and 0.05% (w/v) DDM) by ultrafiltration for 3-times.

Absorption spectra of purified proteins were recorded using a UV-2450 spectrophotometer (Shimadzu, Japan) at room temperature in Buffer A. The retinal composition in PngR was analyzed by high-performance liquid chromatography (HPLC) as described previously^39^. For dark-adaptation, the samples were kept in the dark condition for more than 72 hrs at 4 °C. For light-adaptation, the samples were illuminated for 3 min at 520 ± 10 nm, where the light power was adjusted to approximately 10 mW cm^-2^. The molar compositions of the retinal isomers were calculated from the areas of the peaks in HPLC patterns monitored at 360 nm using the extinction coefficients of retinal oxime isomers as described previously^39^. For pH titration experiments, the samples were suspended in Buffer A. The pH values of the samples were adjusted to the desired acidic values by adding HCl, after which the absorption spectra were measured at each pH value. All measurements were conducted at room temperature (approx. 25 °C) under room light. After the measurements, the reversibility of the spectral changes was checked to confirm that the sample was not denatured during the measurements. The absorption changes at specific wavelengths were plotted against pH values and the plots were fitted to the Henderson–Hasselbalch equation assuming single p*K*_a_ value as previously described^37^.

Transient time-resolved absorption spectra of the purified proteins from 380 to 700 nm at 5 nm intervals were obtained using a homemade computer-controlled flash photolysis system equipped with an Nd: YAG laser as an actinic light source. Using an optical parametric oscillator, the wavelength of the actinic pulse was tuned at 510 nm for PngR. The pulse intensity was adjusted to 2 mJ per pulse. All data were averaged to improve the signal-to-noise ratio (n = 30). All measurements were conducted at 25 °C. For these experiments, the samples were suspended in Buffer A. After the measurements, the reproducibility of the data was checked to confirm that the sample was not denatured during the measurements. To investigate proton uptake and release during the photocycle, we used the pH indicator pyranine (final concentration = 100 μM, Tokyo Chemical Industry Co., Ltd, Japan), which has been extensively used to monitor light-induced pH changes in various rhodopsins. The pH changes in the bulk environment were measured as the absorption changes of pyranine at 450 nm. The absorption changes of pyranine were obtained by subtracting the absorption changes of samples without pyranine from those of samples with pyranine. The experiments using pyranine were performed in an unbuffered solution containing 1 M NaCl and 0.05% (w/v) DDM (pH 7.0) to enhance the signals. The results of 1000-traces were averaged to improve the signal-to-noise ratio.

### Subcellular localization of PngR in the model diatom

The PngR:eGFP recombinant gene, coding the full length of PngR C-terminally tagged with eGFP, was cloned into the expression vector for the model diatom *Phaeodactylum tricornutum*, pPha-NR^40^, by CloneEZ (GenScript) following the manufacturer’s instruction. The plasmid was electroporated into *P. tricornutum* cells with NEPA21 Super Electroporator (NEPAGENE), and transformed cells were selected with a Zeocin-based antibiotic treatment as described previously^17,41^. Selected clones were observed under an Olympus BX51 fluorescent microscope (Olympus) equipped with an Olympus DP72 CCD color camera (Olympus). The nucleus stained with DAPI and chlorophyll autofluorescence from the plastid were observed with a 420 nm filter by 330 to 385 nm excitation. GFP fluorescence was detected with a 510 to 550 nm filter by 470 to 495 nm excitation.

### Quantitative model of C concentration in a diatom: CFM-CC

#### Membrane transport model

In this model, we combined membrane transport of CO_2_ and C fixation (Fig. 4). Parameter definitions, units and values are provided in Supporting Information Table S2 and S3, respectively. The key model equation is the balance of CO_2_ concentration in cytosol [*CO*_*2*_]_*p*_:

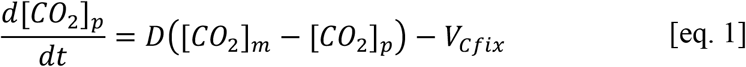

where *t* is time (units), *D* is diffusion coefficient, [*CO*_*2*_]_*m*_ is CO_2_ concentration (units) in the middle space. Here the first term represents the diffusion of CO_2_ from the middle space to cytosol and the second term *V*_*cfix*_ represents the C fixation rate following Michaelis–Menten kinetics^22,42^:

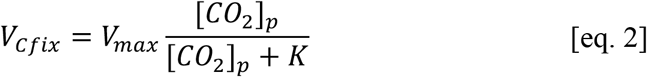

where*V*_*max*_ is maximum CO_2_ fixation rate (units), and *K* is a half saturation constant (units). [*CO*_*2*_]_*m*_ is obtained based on the carbonate chemistry in the middle space (see below). Under the steady-state, [eq. 1] with [eq. 2] becomes the following quadratic relationship for [*CO*_*2*_]_*p*_:

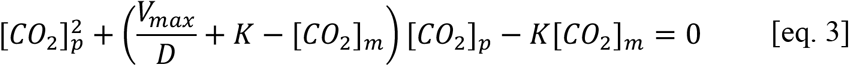

Solving this equation for 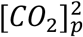 leads to:

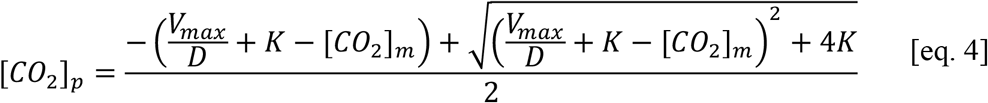

Note that the other solution for the negative rout is unrealistic since it may lead to the overall negative value of [*CO*_*2*_]_*p*_. Once we obtain [*CO*_*2*_]_*p*_, we can then calculate the rate of C fixation *V*_*cfix*_ with [eq. 2].

Also, from [eq. 4], we can get two extreme solutions. First, when *V*_*max*_ ≪ *D* (i.e., when the CO_2_ uptake capacity is small relative to the speed of CO_2_ diffusion) [eq. 4] leads to

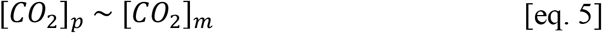

With this relationship and [eq. 2], *V*_*cfix*_ is computed as follows:

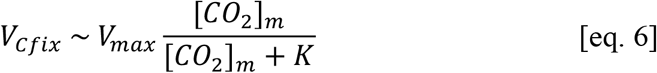

Second, when *V*_*max*_ ≫ *D* (i.e., when the CO_2_ uptake capacity is high relative to the CO_2_ diffusion across the membrane) [eq. 4] becomes

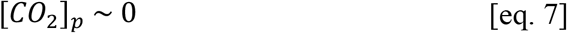

Under the steady state, [eq. 1] becomes

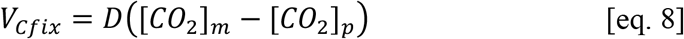

and plugging [eq. 7] into [eq. 8] leads to

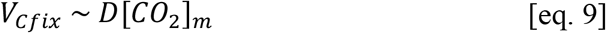

and *V*_*cfix*_ is calculated. We note that [*CO*_*2*_]_*p*_ > [*CO*_*2*_]_*m*_ could occur if there were membrane-bound transporters for HCO_3_^-^located on each membrane between the middle space and plastid^22^. However, such a set of transporters have not been discovered^20^. Thus, our model is fidelity to the current state of knowledge. Even if [*CO*_*2*_]_*p*_ > [*CO*_*2*_]_*m*_ was the case, moderately decreased pH_m_ and thus increased [*CO*_*2*_]_*m*_ could be useful since they would decrease the gradient of CO_2_ across the membranes (i.e., [*CO*_*2*_]_*p*_ vs [*CO*_*2*_]_*m*_), mitigating the diffusive loss of CO_2_ from the plastid.

#### Carbonate chemistry in the middle space

The above equations can be solved once we obtain [*CO*_*2*_]_*m*_. The model uses a given DIC (dissolved inorganic C) concentration in the middle space [*DIC*]_*m*_ to calculate [*CO*_*2*_]_*m*_ following the established equations for carbon chemistry ^43^.

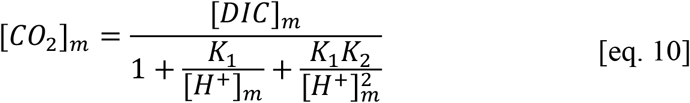

where [*H*^+^]_*m*_ is H^+^ concentration (thus 10^-pH^ mol l^-1^) in the middle space and *K*_1_ and *K*_2_ are temperature and salinity dependent parameters ^43,44^:

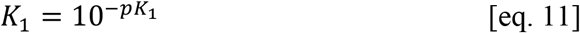

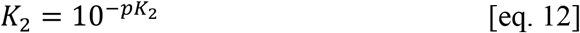

where

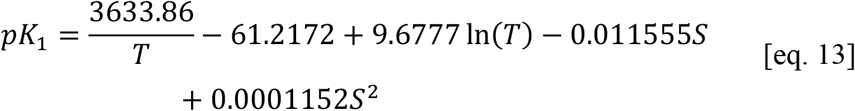

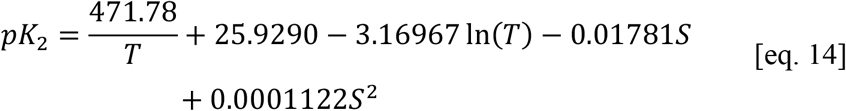

## Code availability

Code for CFM-CC is freely available from GitHub/Zenodo at https://zenodo.org/record/5182712 (DOI: 10.5281/zenodo.5182712).

## Acknowledgements

We thank Yusuke Matsuda for useful discussion. This work was supported by JSPS KAKENHI Grant Numbers 18K19224 and 18H04136 to S.Y., 19H04727, 21H00404 and 21H02446 to Y.S., and 19H03274 to R.K., and NSF grans OPP1745036 to A.M. This research was partially supported by the Interdisciplinary Collaborative Research Program of the Atmosphere and Ocean Research Institute, the University of Tokyo.

## Author contributions

S.Y., A.M., Y.S., and R.K. designed the research; S.Y., T.A., K.K., K.Inomura. M.H., Y.N., M.K., K.Ifuku. and R.K. performed the research; S.Y., K.K., K.Inomura., Y.N., M.K., G.A., H.M.,T.Y., H.F., A.M., Y.S. and R.K. analyzed the data; and S.Y., K.Inomura., Y.S., and R.K. wrote the paper.

## Additional information

Supplementary information is available for this paper.

**Correspondence** and requests for materials should be addressed to S.Y.

