## Supporting Information for "Proton-pumping rhodopsins in marine diatoms"

**This file includes**

Supplementary Results and Discussion p. 2–3

Extended Data Figs. 1–8 p. 4–11

Supplementary Tables S2–S3 p. 12

Supplementary References p. 13

### Supplementary Results and Discussion

#### *Photocycle of PngR*

To analyze the photocycle of PngR, we performed flash-photolysis experiment. Extended Data Fig. 5A shows the flash-induced difference spectra over the spectral range of 380–700 nm. The depletion and recovery of absorbance at ~ 510 nm correspond to the bleaching of the original state, while an increase and decrease of absorbance at ~ 410 and 570 nm were characteristically observed. Extended Data Fig. 5B shows the time courses of the difference absorbance changes at the three wavelengths of 410, 510 and 570 nm. Following the illumination, an absorption increase at ~ 570 nm was observed together with the depletion of the original state. An absorption increase at ~ 410 nm was then observed with a concomitant absorption decrease at ~ 570 nm within 0.5 ms. Considering the temporal and spectral ranges of the absorption changes, the absorbances at 570 and 410 nm were tentatively attributed to the K- and M-intermediates, respectively. The absorbance at ~ 410 nm decreased with the concomitant absorbance increase at ~ 570 nm, which was tentatively assigned as the O-intermediate, within 10 ms. Finally, the absorbance at ~ 570 nm was depleted with recovery of the original state within 300 ms. Thus, after the light absorption, PngR sequentially forms K-, M- and O-intermediates, and then returns to the original state. To estimate the decay time constants of the intermediates, the temporal absorption changes at 410, 510 and 570 nm were fitted with a triple-exponential function assuming the irreversible sequential model. The decay time constants of the K-, M- and O-intermediates were estimated as 0.061, 0.83 and 61 ms, respectively. Finally, we investigated how proton uptake and release happen during the photocycle since PngR exhibits a proton pumping function.

Proton uptake and release by PngR during the photocycle were detected as the time course of the absorbance changes at 450 nm by using pyranine, a pH-sensitive dye. Pyranine works as a pH indicator, and the absorbance of pyranine at 450 nm was decreased under acidic conditions. As a result, the absorbance of pyranine increased within 5 ms and then decreased within 100 ms (Extended Data Fig. 5B), which indicates that the substrate proton was first taken up from the bulk solution and then released from PngR during photocycle. These absorbance changes were observed coinstantaneous with those of O-intermediate, which suggests that proton uptake and release are coincident with the formation and decay of O-intermediate, respectively.

Based on these results, we propose a photocycle model of PngR as shown schematically in Extended Data Fig. 5C.

**Extended Data Figs.**

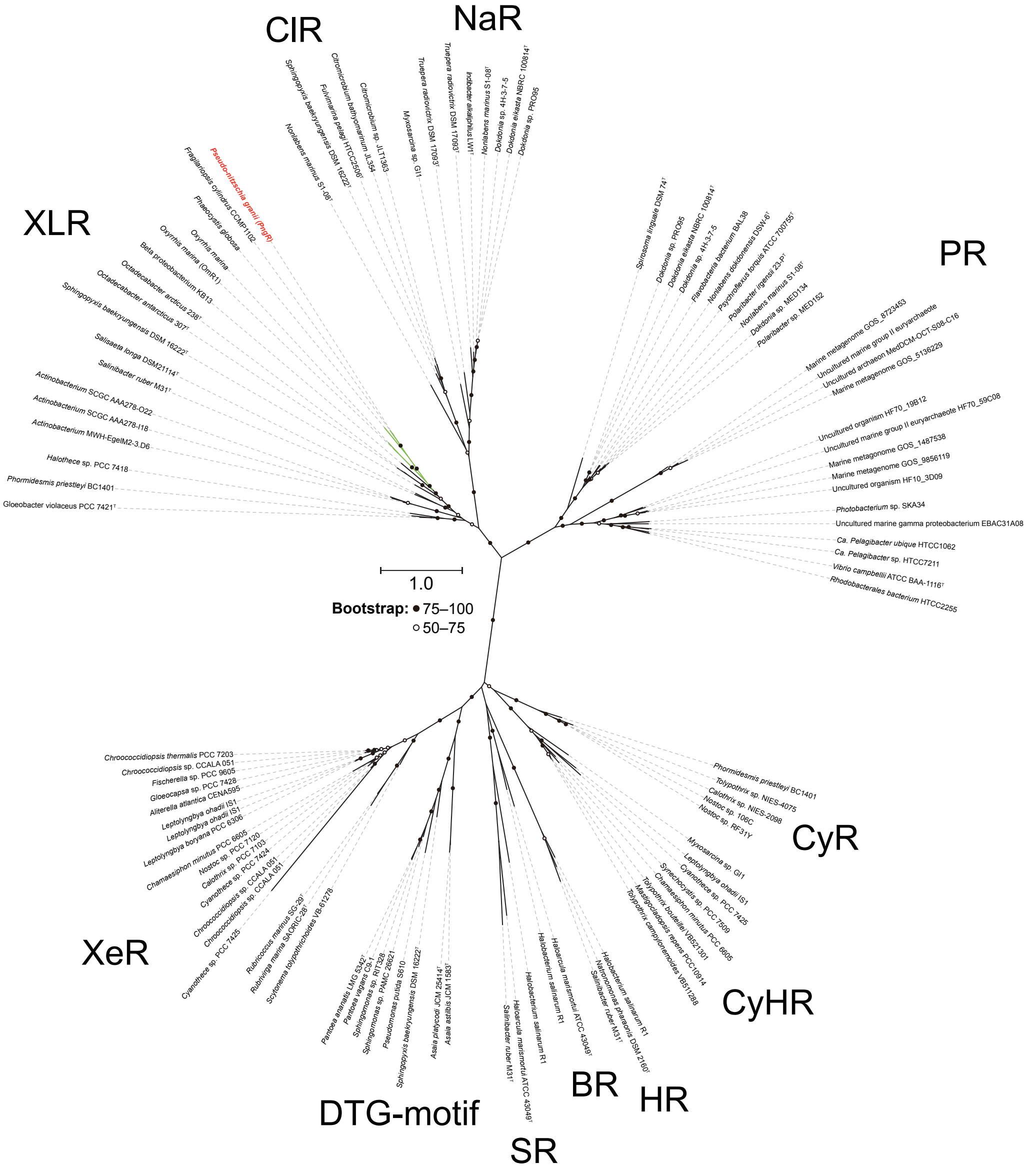


**Extended Data Fig. 1. Phylogenetic position of diatom rhodopsin and the rhodopsin sequences used in this analysis.** A maximum likelihood tree of amino acid sequences of microbial rhodopsins. Diatom rhodopsin (PngR) is indicated by red character and bootstrap probabilities (≥ 50%) are indicated by black and white circles. Green branches indicate eukaryotic rhodopsins used in this analysis, and black branches indicate prokaryotic rhodopsins. Rhodopsin clades are as follows: XLR (Xanthorhodopsin-like rhodopsin), ClR ($Cl^-$-pumping rhodopsin), NaR ($Na^+$-pumping rhodopsin), PR (proteorhodopsin), XeR (xenorhodopsin), DTG-motif rhodopsin, SR (sensory rhodopsin-I and sensory rhodopsin-II), BR (bacteriorhodopsin), HR (halorhodopsin), CyHR (cyanobacterial halorhodopsin), and CyR (cyanorhodopsin).

```
                                                                                       85    89
                                                                                      (91)  (95)
PngR  ---------- ---------- ---------- --MVTANQFT IVYD--VLSF SLATMMATTI FLWMRVPSVH
XR    --MLQEL--- ---------- ---------- -PTLTPGQYS LVFN--MFSF TVATMTASFV FFVLARNNVA
GR    M-LMTVFSSA PELALLGSTF AQVDPSNLSV SDSLTYGQFN LVYN--AFSF AIAAMFASAL FFFSAQALVG
BR    --MLELLPTA V--------- ---------- -EGVSQAQIT GRPEWIWLAL GTALMGLGTL YFLVKGMGVS
PR    MKLLLILGSV IALPTFAAGG ---------- -GDLDASDYT GV----SFWL VTAALLASTV FFFVERDRVS
NaR   --MTQELGNA NFENFIGAT- ---------- -EGFSEIAYQ FTSH--ILTL GYAVMLAGLL YFILTIKNVD

                                                                       85    89
                                                                      (91)  (95)
PngR  -EKYKSALII SGLVTFIAAY HYLRIFNSWT EAYEWTAE-G E--LSATASP FNDAY--RYM DWLLTVPLLL
XR    -PKYRISMMV SALVVFIAGY HYFRITSSWE AAYA-LQN-G M--YQPTGEL FNDAY--RYV DWLLTVPLLT
GR    -QRYRLALLV SAIVVSIAGY HYFRIFNSWD AAYV-LEN-G V--YSLTSEK FNDAY--RYV DWLLTVPLLL
BR    DPDAKKFYAI TTLVPAIAFT MYLSMLLGYG LT-------- ---MVPFGGE QNPIYWARYA DWLFTTPLLL
PR    -AKWKTSLTV SGLVTGIAFW HYMYMRGVWI ET-------- --------GD SPTVF--RYI DWLLTVPLLI
NaR   -KKFQMSNIL SAVVMVSAFL LLYAQAQNWT SSFTFNEEVG RYFLDPSGDL FNNGY--RYL NWLIDVPMLL

 96
(102)
PngR  IEIILVMKLP ADESRSKATT LGIASAAMIA IGYPGELAMS DDSLGTRWVY WIGAMLPFLY IVQTLLVGLN
XR    VELVLVMGLP KNERGPLAAK LGFLAALMIV LGYPGEVSEN AALFGTRGLW GFLSTIPFVW ILYILFTQLG
GR    VETVAVLTLP AKEARPLLIK LTVASVLMIA TGYPGEISDD IT---TRIIW GTVSTIPFAY ILYVLWVELS
BR    LDLALLV--- -DADQGTILA LVGADGIMIG TGLVGALTKV YS---YRFVW WAISTAAMLY ILYVLFFGFT
PR    CEFYLILAAA TNVAGSLFKK LLVGSLVMLV FGYMGEAGIM AA-----WPA FIIGCLAWVY MIYELWAGEG
NaR   FQILFVVSLT TSKFSSVRNQ FWFSGAMMII TGYIGQFYEV SNLT-AFLVW GAISSAFFFH ILWVMKKVIN

                                                                           212   216
                                                                          (227) (231)
PngR  DATQSEADPA IRDLIKGVQW WTVISWCTYP VVYVFPMM-G ITG-----TN AIVGIQ---L GYSVSDIISK
XR    DTIQRQS-SR VSTLLGNARL LLLATWGFYP IAYMIPMA-F PEAFPSNTPG TIVALQ---V GYTIADVLAK
GR    RSLVRQP-AA VQTLVRNMRW LLLLSWGVYP IAYLLPML-G VSG-----TS AAVGVQ---V GYTIADVLAK
BR    SKAESMR-PE VASTFKVLRN VTVVLWSAYP VVWLIGS--- -EG------A GIVPLNIETL LFMVLDVSAK
PR    KSACNTASPA VQSAYNTMMY IIIFGWAIYP VGYFTGYL-M GDG------G SALNLN---L IYNLADFVNK
NaR   EGKEGIS-PA GQKILSNIWI LFLISWTLYP GAYLMPYLTG VDGFLYS-ED GVMARQ---L VYTIADVSSK

PngR  CGVGLMIYQI TIAKSYAL-- --KKGNEETP LYG------- -----
XR    AGYGVLIYNI AKAKSEEEGF NVSEMVEPAT ASA------- -----
GR    PVFGLLVFAI ALVKTKAD-- --QESSEPHA AIGAAANKSG GSLIS
BR    VGFGLILL-- ---RSRAI-F GEAEAPEPSA GDGAAATSD- -----
PR    ILFGLIIWNV AVKESSNA-- ---------- ---------- -----
NaR   VIYGVLLGNL AITLSKNKEL --VEANS--- ---------- -----
```

**Extended Data Fig. 2. Sequence alignment of rhodopsins.** The accessions and rhodopsin families are as follows: PngR (AJA37445.1, XLR), XR (WP_011404249.1, XLR), GR (BAC88139.1, XLR), BR (CAP14056.1, BR), PR (AAG10475.1, PR), and NaR (BAN14808.1, NaR). All rhodopsins except NaR function as proton pump. Columns of functionally important residues are shown in bold. The numbers above the columns indicate amino acid numbers in BR, and PngR in parentheses. Known functions are as follows: primary proton acceptor (Asp85 in BR), proton donor (Glu96), counterion (Asp212), Schiff base (Lys216). Two carboxylates, Asp (D) and Glu (E), are shown in blue, and Schiff base Lys (K) is shown in red.

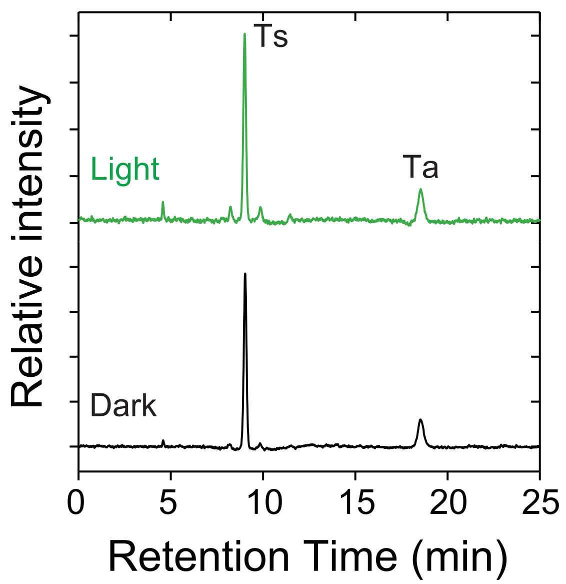


**Extended Data Fig. 3. Retinal configuration of PngR.** HPLC patterns of retinal isomers of PngR with (green line) and without (black line). Ts and Ta represent all-*trans*-15-*syn* and all-*trans*-15-*anti* retinal oximes, respectively.

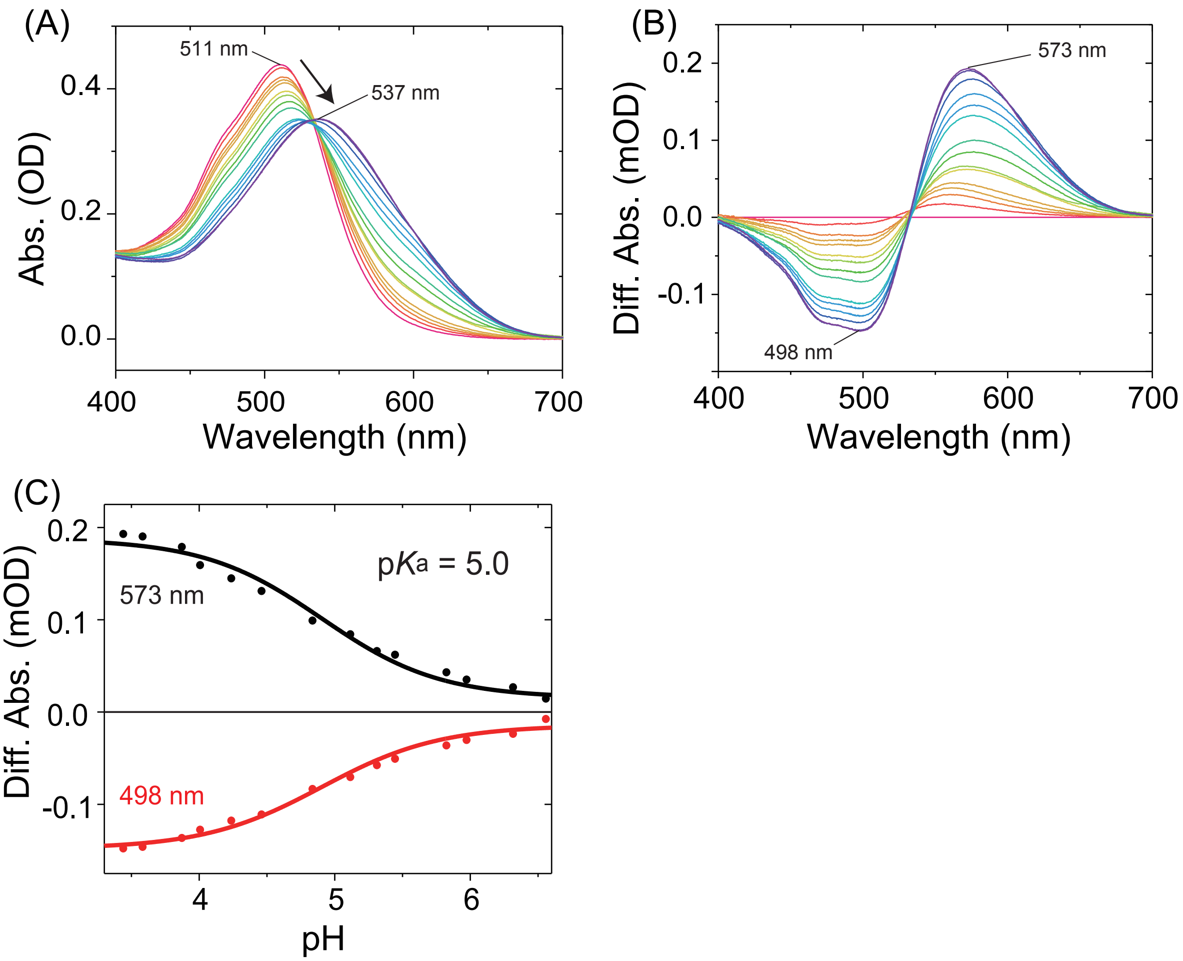


**Extended Data Fig. 4. pH-induced spectral changes of PngR.** (A) Absorption spectra of PngR at acidic pH from 7.0 to 3.4 in Buffer A containing 50 mM Tris–HCl, 1 M NaCl and 0.05% (w/v) DDM. (B) Difference absorption spectra; each spectrum was obtained by subtracting the spectrum at pH 7.0. (C) Plots of the difference absorbance at 498 and 573 nm against the pH values. The titration curve was analyzed using the Henderson–Hasselbalch equation assuming single pKa value (solid lines).

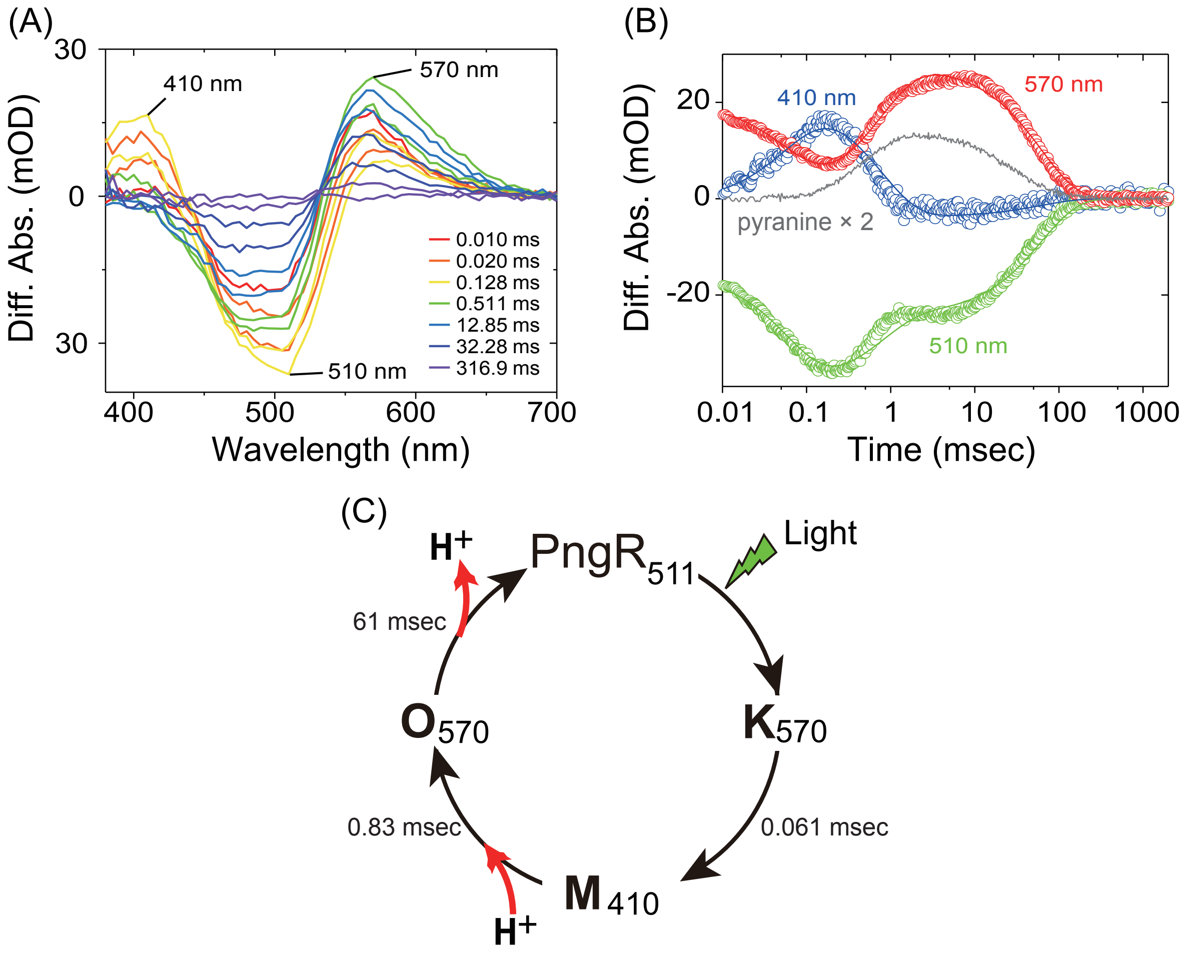


**Extended Data Fig. 5. Photocycle and proton transport model of PngR.** (A) Flash-induced difference absorption spectra over the spectral range of 380 to 700 nm in Buffer A containing 50 mM Tris-HCl (pH 7.0), 1 M NaCl and 0.05% (w/v) DDM. (B) Time courses of absorbance changes at 410, 510, and 570 nm. The black solid lines indicate the fitting curves. The absorption changes of pyranine monitored at 450 nm were enlarged 2 times and are shown as a gray solid line. (C) Proposed photo cycle model of PngR with the timing of the proton release and uptake.

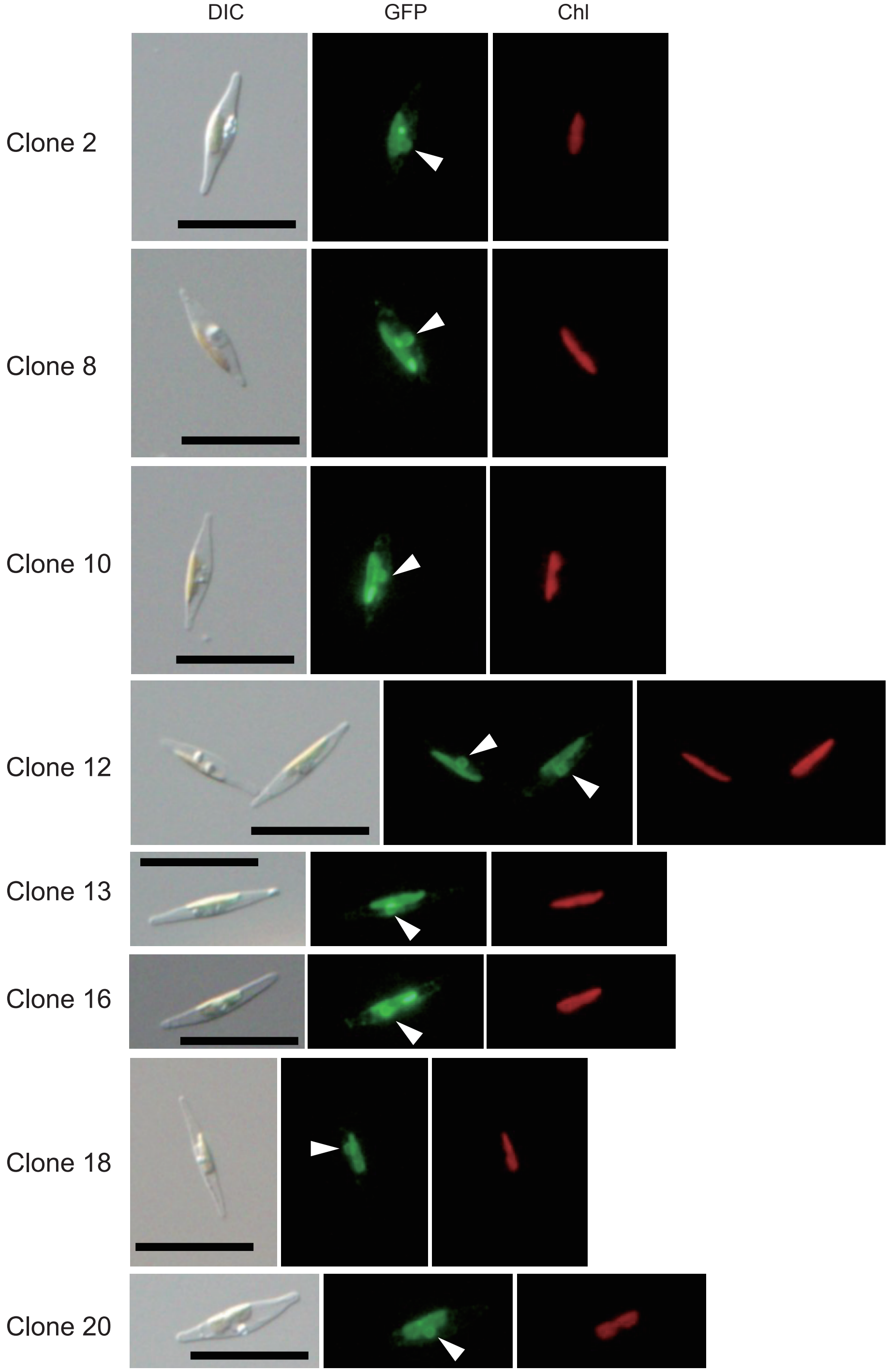


**Extended Data Fig. 6. Subcellular localization of the exogenously introduced PngR and plastid in diatom cells.** Transformed diatom cells were observed with DIC (Differential Interface Contrast) (Left). Green fluorescence from the recombinant protein (GFP) and the chlorophyll autofluorescence (Chl) are shown in center and right, respectively. The triangles show the location of nucleus, and GFP surrounds the nucleus. Scale bar indicates 20 µm.

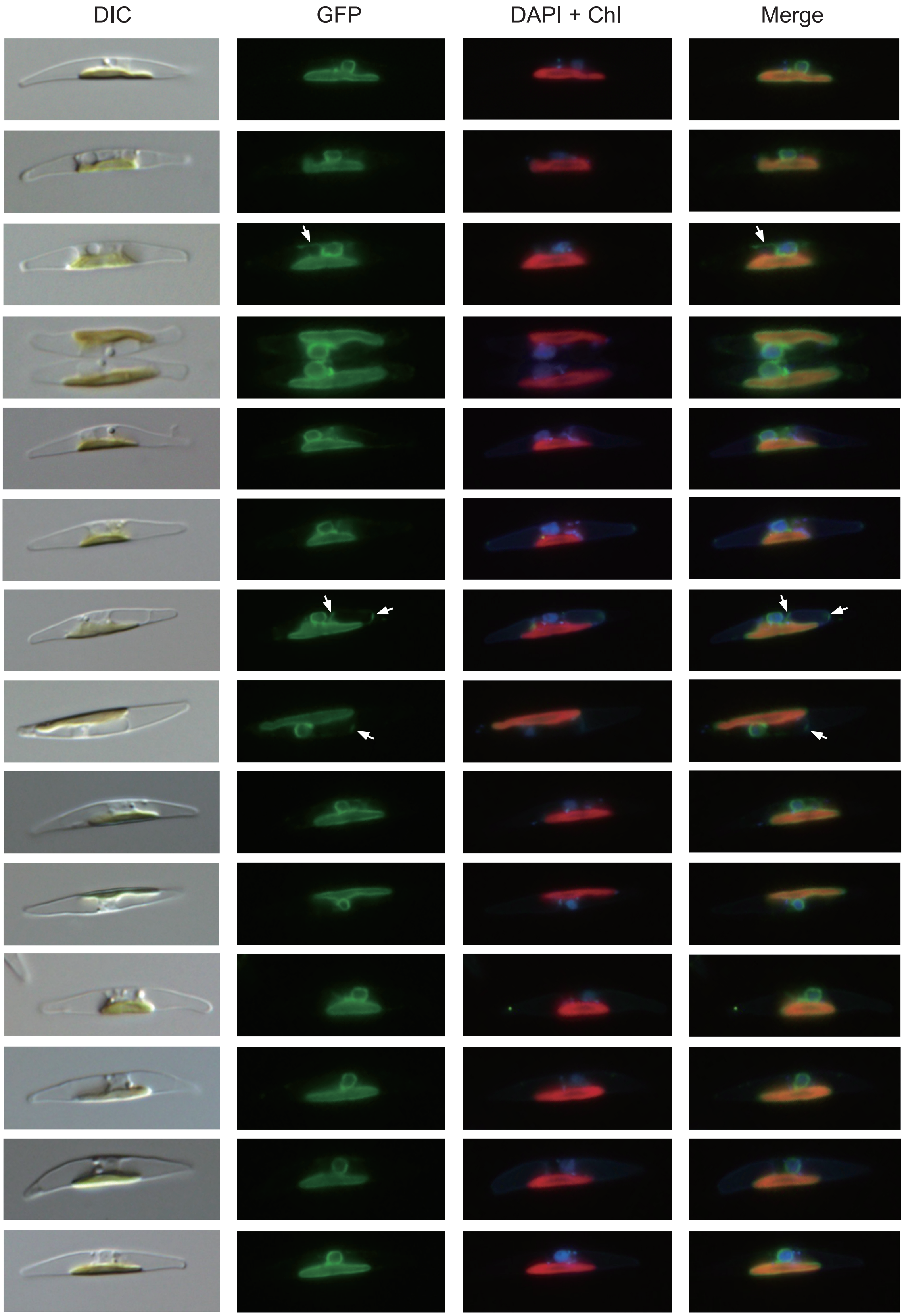


96

**Extended Data Fig. 7. Subcellular localization of the exogenously introduced PngR, nucleus and plastid in diatom cells.** Transformed diatom cells were observed with DIC (Differential Interface Contrast) (Left). Green fluorescence from the recombinant protein (GFP) (Left center). The nuclear DNA stained with DAPI and the chlorophyll autofluorescence (DAPI + Chl) and merged image (Merge) are shown in right center and right, respectively. Arrows indicate GFP fluorescence outside the chloroplast; most of the GFP fluorescence is localized to the CERM, but it may also be observed in other organelles (e.g., vacuoles and periplasmic membranes).

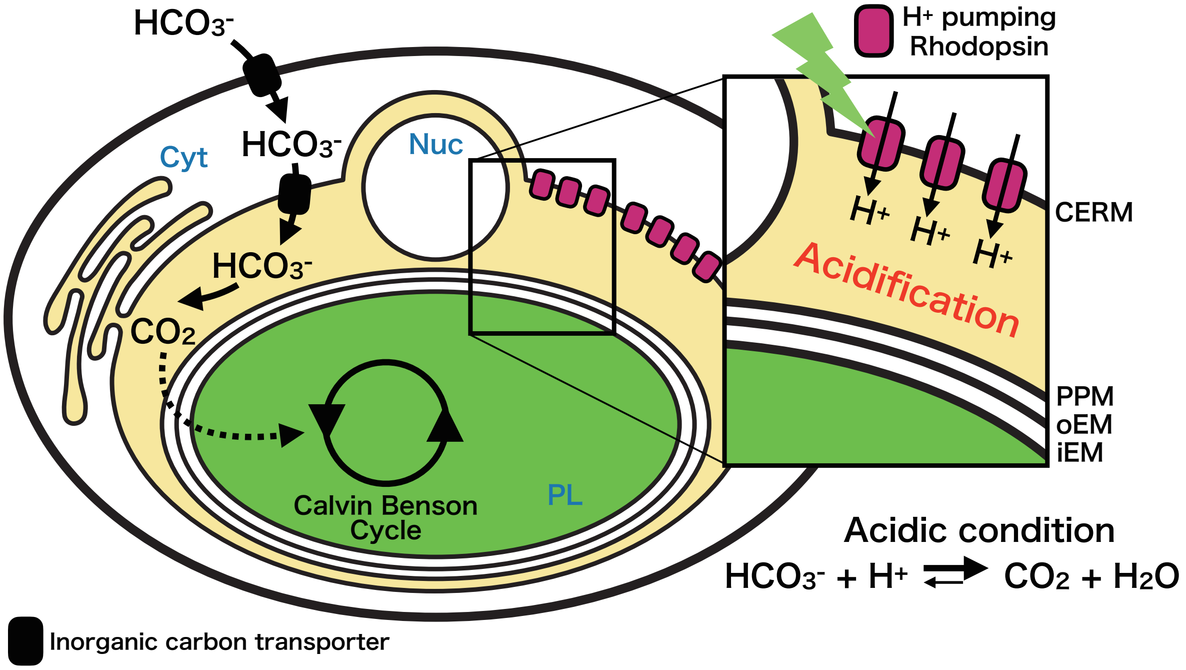


**Extended Data Fig. 8. A proposed model that proton transport by rhodopsin is involved in CCM.** The proton transport of rhodopsin acidifies the region (the middle space) surrounded by the membrane of CERM and PPM. Abbreviation are as follows: Cyt (Cytosol), Nuc (Nucleus), PL (Plastid), CERM (Chloroplast endoplasmic reticulum membrane), PPM (Periplastidial membrane), oEM (Outer plastid envelop membrane) and iEM (Internal plastid envelop membrane).                    .

### Supplementary Tables

Table S2 Parameters, units and definitions. Parameters are listed roughly in the order of appearance.

| Parameter | Unit | Definition |
|---|---|---|
| $[CO_2]_p$ | µmol L$^{-1}$ | $CO_2$ concentration in the plastid |
| $t$ | d | time |
| $D$ | d$^{-1}$ | diffusion coefficient |
| $[CO_2]_m$ | µmol L$^{-1}$ | $CO_2$ concentration in the middle space |
| $V_{max}$ | µmol L$^{-1}$ d$^{-1}$ | maximum C fixation rate |
| $K$ | µmol L$^{-1}$ | Half-saturation constant for C fixation |
| $V_{Cfix}$ | µmol L$^{-1}$ d$^{-1}$ | C fixation rate |
| $[DIC]_m$ | µmol L$^{-1}$ | DIC concentration in the middle space |
| $[H^+]_m$ | mol kg$^{-1}$ | $H^+$ concentration in the middle space |
| $K_1$ | mol kg$^{-1}$ | temperature and salinity dependent constant 1 |
| $K_2$ | mol kg$^{-1}$ | temperature and salinity dependent constant 2 |
| $T$ | K | temperature |
| $S$ | ‰ | salinity |

Table S3 Parameter values

| Parameter | Value | Unit |
|---|---|---|
| $K$ | 44*1 | µmol L$^{-1}$ |
| $[DIC]_m$ | 993*2 | µmol L$^{-1}$ |
| $T$ | 298.15*3 | K |
| $S$ | 35*4 | ‰ |

*1 Based on the middle value from [1]

*2 Observed value of DIC in a diatom *Phaeodactylum tricornutum*[2]. This is the only value we found for diatoms.

*3 Typical room temperature.

*4 Typical values of seawater (note that $K_1$ and $K_2$ values are relatively insensitive to $S$ values).
